# Efficient DNA-based data storage using shortmer combinatorial encoding

**DOI:** 10.1101/2021.08.01.454622

**Authors:** Inbal Preuss, Michael Rosenberg, Zohar Yakhini, Leon Anavy

## Abstract

With the world generating digital data at an exponential rate, DNA has emerged as a promising archival medium. It offers a more efficient and long-lasting digital storage solution due to its durability, physical density, and high information capacity. Research in the field includes the development of encoding schemes, which are compatible with existing DNA synthesis and sequencing technologies. Recent studies suggest leveraging the inherent information redundancy of these technologies by using composite DNA alphabets. A major challenge in this approach involves the noisy inference process, which prevented the use of large composite alphabets. This paper introduces a novel approach for DNA-based data storage, offering a 6.5-fold increase in logical density over standard DNA-based storage systems, with near zero reconstruction error. Combinatorial DNA encoding uses a set of clearly distinguishable DNA shortmers to construct large combinatorial alphabets, where each letter represents a subset of shortmers. The nature of these combinatorial alphabets minimizes mix-up errors, while also ensuring the robustness of the system.

As this paper will show, we formally define various combinatorial encoding schemes and investigate their theoretical properties, such as information density, reconstruction probabilities and required synthesis, and sequencing multiplicities. We then suggest an end-to-end design for a combinatorial DNA-based data storage system, including encoding schemes, two-dimensional error correction codes, and reconstruction algorithms. Using *in silico* simulations, we demonstrate our suggested approach and evaluate different combinatorial alphabets for encoding 10KB messages under different error regimes. The simulations reveal vital insights, including the relative manageability of nucleotide substitution errors over shortmer-level insertions and deletions. Sequencing coverage was found to be a key factor affecting the system performance, and the use of two-dimensional Reed-Solomon (RS) error correction has significantly improved reconstruction rates. Our experimental proof-of-concept validates the feasibility of our approach, by constructing two combinatorial sequences using Gibson assembly imitating a 4-cycle combinatorial synthesis process. We confirmed the successful reconstruction, and established the robustness of our approach for different error types. Subsampling experiments supported the important role of sampling rate and its effect on the overall performance.

Our work demonstrates the potential of combinatorial shortmer encoding for DNA-based data storage, while raising theoretical research questions and technical challenges. These include the development of error correction codes for combinatorial DNA, the exploration of optimal sampling rates, and the advancement of DNA synthesis technologies that support combinatorial synthesis. Combining combinatorial principles with error-correcting strategies paves the way for efficient, error-resilient DNA-based storage solutions.

## 2 Introduction

DNA is a promising media storage candidate for long-term data archiving, due to its high information density, long-term stability, and robustness. In recent years, several studies have demonstrated the use of synthetic DNA for storing digital information on a megabyte scale, exceeding the physical density of current magnetic tape-based systems by roughly six orders of magnitude [1] [2].

Research efforts in the field of DNA-based storage systems have mainly focused on the application of various encoding schemes, while relying on standard DNA synthesis and sequencing technologies. These include the development of error correcting codes for the unique information channel of DNA-based data storage [3] [4] [5] [6] [7]. Random access capabilities for reading specific information stored in DNA also require advanced coding schemes [8] [9] [10]. Yet, despite the enormous benefits potentially associated with capacity, robustness, and size, existing DNA-based storage technologies are characterized by inherent information redundancy. This is due to the nature of DNA synthesis and sequencing methodologies, which process multiple molecules that represent the same information bits in parallel. Recent studies suggest exploiting this redundancy to increase the logical density of the system, by extending the standard DNA alphabet using composite letters (also referred to as degenerate bases), and thereby encoding more than 2 bits per letter [11] [12].

In this approach, a composite DNA letter uses all four DNA bases (A, C, G, and T), combined or mixed in a specified predetermined ratio *σ* = (*σ*_*A*_, *σ*_*C*_, *σ*_*G*_, *σ*_*T*_). A resolution parameter *k* = *σ*_*A*_ + *σ*_*C*_ + *σ*_*G*_ + *σ*_*T*_ is defined, for controlling the alphabet size. The full composite alphabet of resolution *k*, denoted *ϕ*_*k*_, is the set of all composite letters, so that *∑*_*i*∈(*A,C,G,T*}_*σ*_*i*_ = *k*. Writing a composite letter is done by using a mixture of the DNA bases determined by the letter’s ratio in the DNA synthesis cycle. Current synthesis technologies produce multiple copies, and by using the predetermined base mixture each copy will contain a different base, thus preserving the ratio of the bases at the sequence population level.

While the use of numerical ratios supports higher logical density in composite synthesis, it also introduces challenges related to the synthesis and inference of exact ratios. Combinatorial approaches, which also consist of mixtures, address these challenges in a different way. Studies by Roquet et al. (2021) and Yan et al. (2023) contribute significantly to advancing DNA-based data storage technology. To encode and store data, Roquet et al. focus on a novel combinatorial assembly method for DNA. Yan et al. extend the frontiers of this technology by enhancing the logical density of DNA storage, using enzymatically-ligated composite motifs [13] [14].

In this paper, we present a novel approach for encoding information in DNA, using combinatorial encoding and shortmer DNA synthesis. The method described herein leverages the advantages of combinatorial encoding schemes, while relying on existing DNA chemical synthesis methods with some modifications. Using shortmer DNA synthesis also minimizes the effect of synthesis and sequencing errors. We formally define shortmer-based combinatorial encoding schemes, explore different designs, and analyze their performance. We use computer-based simulations of an end-to-end DNA-based data storage system built on combinatorial shortmer encodings, and study its performance. To demonstrate the potential of our suggested approach and experimentally test its validity, we performed an assembly-based molecular implementation of the proposed combinatorial encoding scheme, and analyzed the resulting data. Finally, we discuss the potential of combinatorial encoding schemes and the future work required to enable these schemes in large-scale DNA-based data storage systems and other DNA data applications. All the code and data used in this study are freely available at:

https://github.com/InbalPreuss/dna_storage_shortmer_simulation

https://github.com/InbalPreuss/dna_storage_experiment

The raw data is available in ENA (European Nucleotide Archive).

The datasets generated and/or analysed during the current study are available in ENA (European Nucleotide Archive) the repository, Accession Number - ERR12364864

## 3 Results

### 3.1 Shortmer combinatorial encoding for DNA storage

We suggest a novel method to extend the DNA alphabet while ensuring near-zero error rates. Let *Ω* be a set of DNA k-mers that will serve as building blocks for our encoding scheme. Denote the elements in *Ω* as *X*_1_, …, *X*_*N*_. Elements in *Ω* are designed to be sufficiently different from each other, to minimize mix-up error probability. Formally, the set is designed to satisfy *d*(*X*_*i*_, *X*_*j*_) ≥ *d*; ∀ *i* ≠ *j*, with the minimal Hamming distance *d* serving as a tunable parameter. Note that *N* = |*Ω*| ≤ 4^*k*^. The elements in *Ω* will be used as building blocks for combinatorial DNA synthesis in a method similar to the one used for composite DNA synthesis [12]. Examples of k-mer sets *Ω* are presented in Supplementary Section 7.3.

We define a combinatorial alphabet *∑* over *Ω* as follows. Each letter in the alphabet represents a non-empty subset of the elements in *Ω*. Formally, a letter *σ* ∈ *∑*, representing a subset *S* ⊆ *Ω*/∅, can be written as an N-dimensional binary vector where the indices for which *σ*_*i*_ = 1 represents the k-mers from *Ω* included in the subset S. We denote the k-mers in *S* as *member k-mers* of the letter *σ*. For example, *σ* = (0,1,0,1,1,0) represents *S* = {*X*_2_, *X*_4_, *X*_5_} and |*Ω*| = *N* = 6. Fig. 1a and Fig. 1b illustrates an example of a combinatorial alphabet using *N* = 16, in which every letter represents a subset of size 5 of Ω. Section 3.2 includes a description of the construction of different combinatorial alphabets.

**Fig. 1:**
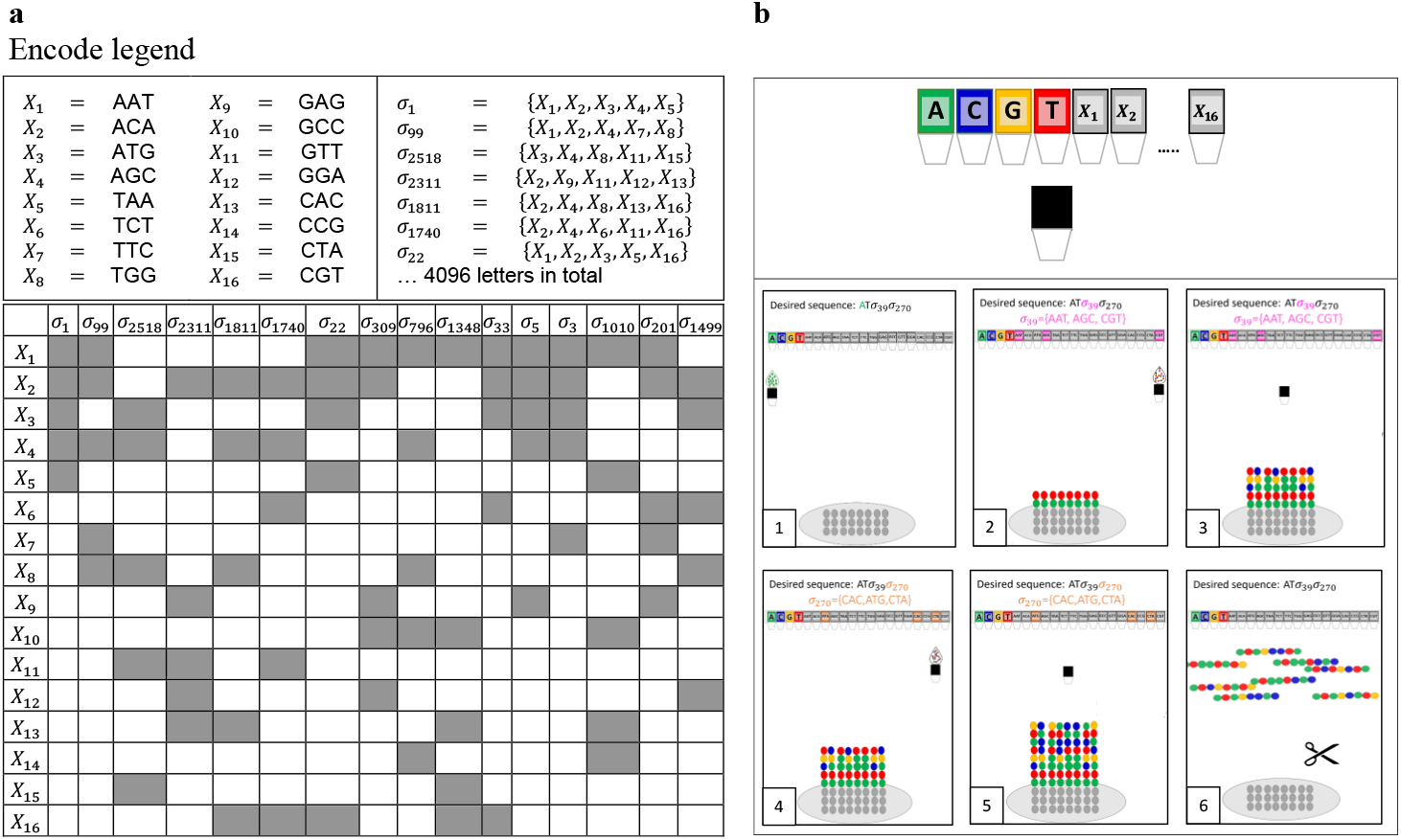
Our combinatorial encoding and synthesis approach. **a**, Schematic view of a combinatorial alphabet (Encode legend). A set of 16 trimers, ***X***_**1**_, …, ***X***_**16**_, is used to construct 4096 combinatorial letters, each representing a subset of 5 trimers as indicated on the right and depicted in the grayed-out cells of the table. **b**, A suggested approach for combinatorial shortmer synthesis. A modified synthesizer would include designated containers for the 16 trimer building blocks and a mixing chamber. Standard DNA synthesis is used for the barcode sequence (1), while the combinatorial synthesis proceeds as follows: The trimers included in the synthesized combinatorial letter are injected into the mixing chamber and introduced into the elongating molecules (2-3). The process repeats for the next combinatorial letter (4-5), and finally, the resulting molecules are cleaved and collected (6).

To write a combinatorial letter *σ* in a specific position, a mixture of the member k-mers of *σ* is synthesized. To infer a combinatorial letter *σ*, a set of reads needs to be analyzed to determine which k-mers are observed in the analyzed position (See Section 3.2 and Section 3.3 for more details). This set of k-mers observed in the sequencing readout and used for inferring *σ* is referred to as *inferred member k-mers*.

From a hardware/chemistry perspective, the combinatorial shortmer encoding scheme is potentially based on using the standard phosphonamidite chemistry synthesis technology, with some alterations (See Fig. 1b, and Supplementary Section 7.1) [15] [16]. First, DNA k-mers are used as building blocks for the synthesis [17]. Such reagents are commercially available for DNA trimers and were used, for example, for the synthesis of codon optimization DNA libraries [18] [19]. In addition, a mixing step will be added to each cycle of the DNA synthesis. Initially, all the member k-mers are added to a designated mixing chamber, and only then is the mixture introduced (for example, by injection) to the elongating molecules. Such systems are yet to be developed.

Similar to composite DNA encoding, combinatorial encoding requires the barcoding of the sequences using unique barcodes composed of standard DNA barcodes. This design enables direct grouping of reads pertaining to the same combinatorial sequence. These groups of reads are the input for the process of reconstructing the combinatorial letters.

The extended combinatorial alphabets allow for a higher logical density of the DNA-based storage system, while the binary nature of the alphabet minimizes error rates.

### 3.2 Binary and binomial combinatorial alphabets

The main parameter that defines a combinatorial encoding scheme is the alphabet *∑*. More specifically, it is the set of valid subsets of *Ω* that can be used as letters. We define two general approaches for the construction of *∑*. Namely, the *binomial encoding* and the *full binary encoding*.

In the *binomial encoding* scheme, only subsets of *Ω* of size exactly *K* represent valid letters in *∑*, so that every letter *σ* ∈ *∑* consists of exactly *K* member k-mers. Therefore, all the letters in the alphabet have the same Hamming weight *K. w*(*σ*) = *K*, ∀*σ* ∈ *∑*. This yields an effective alphabet of size 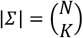 letters, where each combinatorial letter encodes 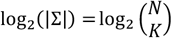 bits. An r-bit binary message requires 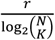 synthesis cycles (and a DNA molecular segment with length 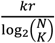 . In practice, we would prefer working with alphabet sizes that are powers of two, where each letter will encode for 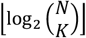 bits. Note that this calculation ignores error correction redundancy, random access primers, and barcodes, which are all required for message reconstruction. See Supplementary Section 7.2 and Fig. 1a, which illustrate a trimer-based binomial alphabet with *N* = 16 and *K* = 5 resulting in an alphabet of size 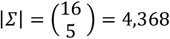 that allows to encode ⌊*log*_2_ (4368)⌋ = 12 bits per letter or synthesis position.

In the *full binary encoding* scheme, all possible nonempty subsets of *Ω* represent valid letters in the alphabet. This yields an effective alphabet of size |*∑*| = 2^*N*^ − 1 letters, each encoding for ⌊*log*_2_(|*∑*|)⌋ = *N* − 1 bits.

From this point on, we focus on the binomial encoding.

### 3.3 Reconstruction probabilities for binomial encoding

In this section, the performance characteristics of binomial encoding is investigated. Specifically, we present a mathematical analysis of the probability of successfully reconstructing the intended message. In Section 3.4 and Section 3.5 results are presented from our simulations and from a small-scale molecular implementation of the binomial encoding, respectively.

#### 3.3.1 Reconstruction of a single combinatorial letter

Since every letter *σ* ∈ *∑* consists exactly of the *K* member k-mers, the required number of reads for observing at least one read of each member k-mer in a single letter follows the coupon collector distribution [20]. The number of reads required to achieve this goal can be described as a random variable 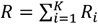, where *R*_1_ = 1 and 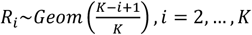 . Hence, the expected number of required reads, is:

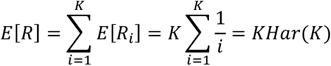

where *Har*(⋅) is the harmonic number.

The expected number of reads required for reconstructing a single combinatorial letter thus remains reasonable for the relevant values of *K*. For example, when using a binomial encoding with *K* = 5 the expected number of reads required for reconstructing a single combinatorial letter is roughly 11.5, which is very close to the experimental results presented in Section 3.5. By Chebyshev’s inequality (See Section 5.1), we can derive a (loose) upper bound on the probability of requiring more than *E*[*R*] + *cK* reads to observe at least one read of each member k-mer, where *c* > 1 is a parameter:

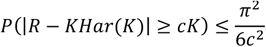

For example, when using a binomial encoding with *K* = 5, the probability of requiring more than 26.5 reads (corresponding to *c* = 3) is bounded by 0.18, which is consistent with the experimental result shown in Fig. 4d.

#### 3.3.2 Reconstruction of a combinatorial sequence

When we examine an entire *K*-subset binomial encoded combinatorial sequence of length *l*, we denote by *R*(*l*) the required number of reads to observe *K* distinct k-mers in every position. Assuming independence between different positions and not taking errors into account, we get the following relationship between *c* and any desired confidence level 1 − *δ* (See Section 5.1 for details):

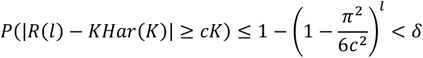

And therefore:

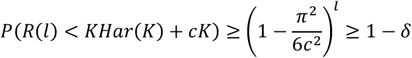

The number of reads required to guarantee reconstruction of a binomial encoded message, at a 1 − *δ* probability, with *K* = 5, and *l* synthesized positions, is thus *KHar*(*K*) + *cK* when 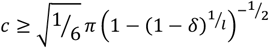.

Table 4 shows several examples of this upper bound. As demonstrated in the simulations and the experimental results, this bound is not tight (See Section 3.4 and Section 3.5).

Note that with an online sequencing technology (such as nanopore sequencing) the sequencing reaction can be stopped after *K* distinct k-mers are confidently observed.

To take into account the probability of observing a k-mer that is not included in Ω (e.g., due to synthesis or sequencing error), we can require that at least *t* > 1 reads of each of the *K* distinct k-mers will be observed. This is experimentally examined in Section 3.5, while the formal derivation of the number of required reads is not as trivial, and will be addressed in future work. The above analysis is based only on oligo recovery, which depends solely on the sampling rate, ignoring possible mix-up errors (i.e., incorrect k-mer readings). This assumption is based on the near-zero mix-up probability attained by the construction of *Ω*, which maximizes the minimal Hamming distance between elements in *Ω*. In Section 3.5, this analysis is compared to experimental results obtained from using synthetic combinatorial DNA.

### 3.4 An end-to-end combinatorial shortmer storage system

We suggest a complete end-to-end workflow for DNA-based data storage with the combinatorial shortmer encoding presented in Fig. 2. The workflow begins with encoding, followed by DNA synthesis, storage, and sequencing, and culminates in a final decoding step. A two-dimensional (2D) error correction scheme, which corrects errors in the letter reconstruction (for example., due to synthesis, sequencing, and sampling errors) and any missing sequences (such as dropout errors), ensures the integrity of the system. Table 1 shows the encoding capacities of the proposed system, calculated on a 1GB input file with standard encoding and three different binomial alphabets. All calculations are based on error correction parameters similar to those previously described (See Section 5.4) [3] [12]. With these different alphabets, up to 6.5-fold increase in information capacity is achieved per synthesis cycle, compared to standard DNA-based data storage.

**Table 1:** Logical densities for selected encoding schemes. The numbers represent encoding a 1 GB binary message using oligos with 14nt barcodes +2nt RS (standard DNA), and 120 payload letters (from **∑**) with 14 extra RS for the payload (the payload and its RS is combinatorial with N and K as indicated).

| Type | $N$ | $K$ | $\binom{N}{K}$ | Bits per letter | Alphabet size | Bits per sequence | Number of sequences | Reed Solomon (RS) | Bits per synthesis cycle, payload only | Bits per synthesis cycle | Fold increase |
| --- | --- | --- | --- | --- | --- | --- | --- | --- | --- | --- | --- |
| Standard |  |  |  | 2 | 4 | 240 | 33,333,334 | 38,095,248 | 1.57 | 1.40 | 1.0 |
| Binomial | 16 | 3 | 560 | 9 | 512 | 1,080 | 7,407,408 | 8,465,616 | 7.05 | 6.30 | 4.5 |
| Binomial | 16 | 5 | 4,368 | 12 | 4,096 | 1,440 | 5,555,556 | 6,349,248 | 9.40 | 8.40 | 6.0 |
| Binomial | 16 | 7 | 11,440 | 13 | 8,192 | 1,560 | 5,128,206 | 5,860,848 | 10.19 | 9.10 | 6.5 |

**Fig. 2:**
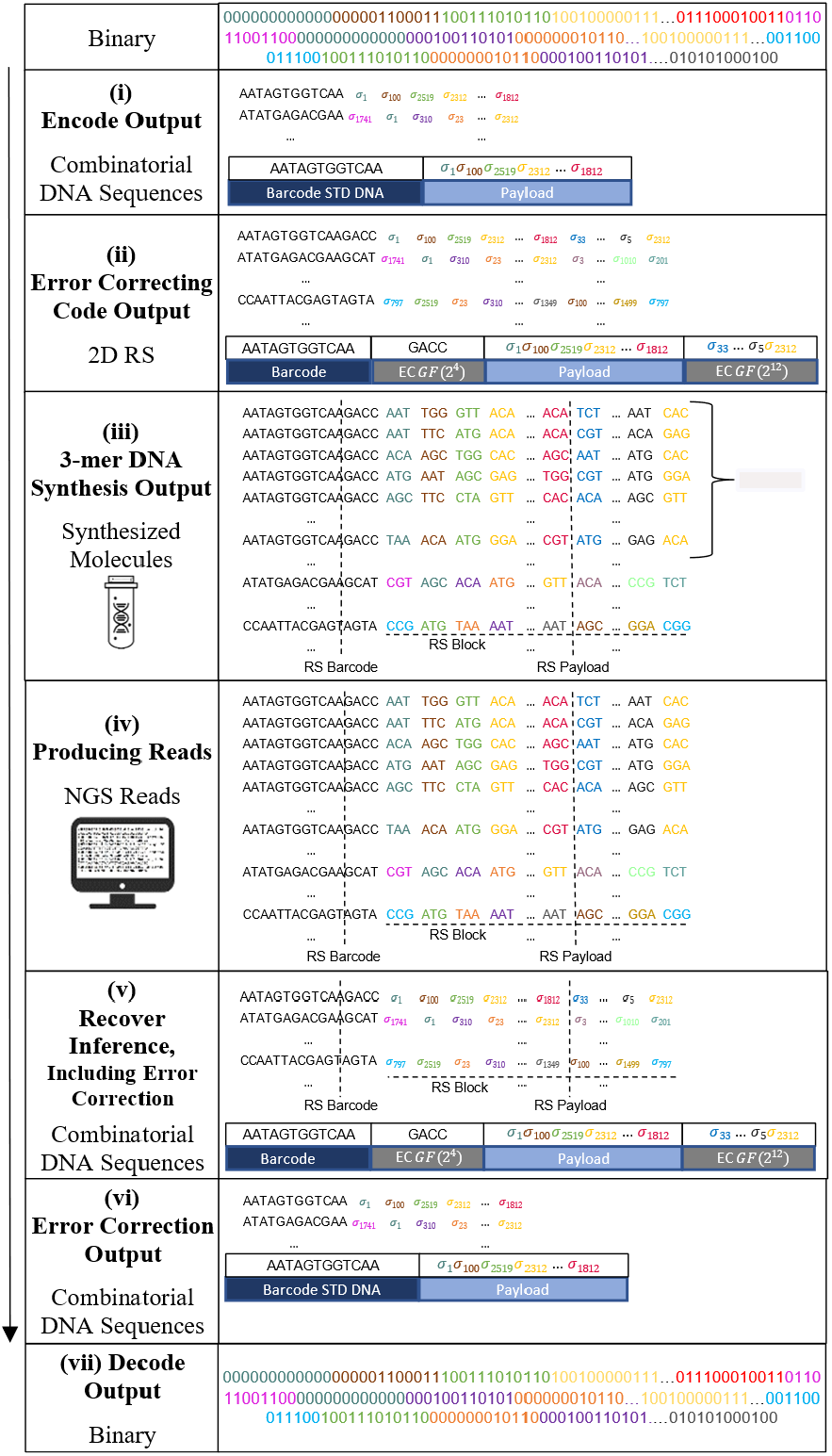
End-to-end workflow of a combinatorial DNA storage system. A binary message is broken into chunks, barcoded, and encoded into a combinatorial alphabet (i). RS encoding is added to each chunk and each column (ii). The combinatorial message is synthesized using combinatorial shormer synthesis (iii) and the DNA is sequenced (iv). Next, the combinatorial letters are reconstructed (v). Finally the message goes through 2D RS decoding (vi), followed by its translation back into the binary message (vii).

An example of the proposed approach, using a binomial alphabet with *N* = 16 and *K* = 5 and two-dimensional Reed Solomon (RS), is detailed below. A binary message is encoded into a combinatorial message using the 4096-letter alphabet. Next, the message is broken into 120 letter chunks, and each chunk is barcoded. The 12nt barcodes are encoded using RS(6,8) over *GF*(2^4^), resulting in 16nt barcodes. Each chunk of 120 combinatorial letters is encoded using RS(120,134) over *GF*(2^12^). Every block of 42 sequences is then encoded using RS(42,48) over *GF*(2^12^) (see Section 5.2 for details).

To better characterize the potential of this proposed system, we implemented an end-to-end simulation using the parameters mentioned above. We simulated the encoding and decoding of 10KB messages with different binomial alphabets and error probabilities, and then measured the resulting reconstruction and decoding rates throughout the process. Fig. 3a depicts a schematic representation of our simulation workflow and indicates how the error rates are calculated (See Section 5.2.4).

**Fig. 3:**
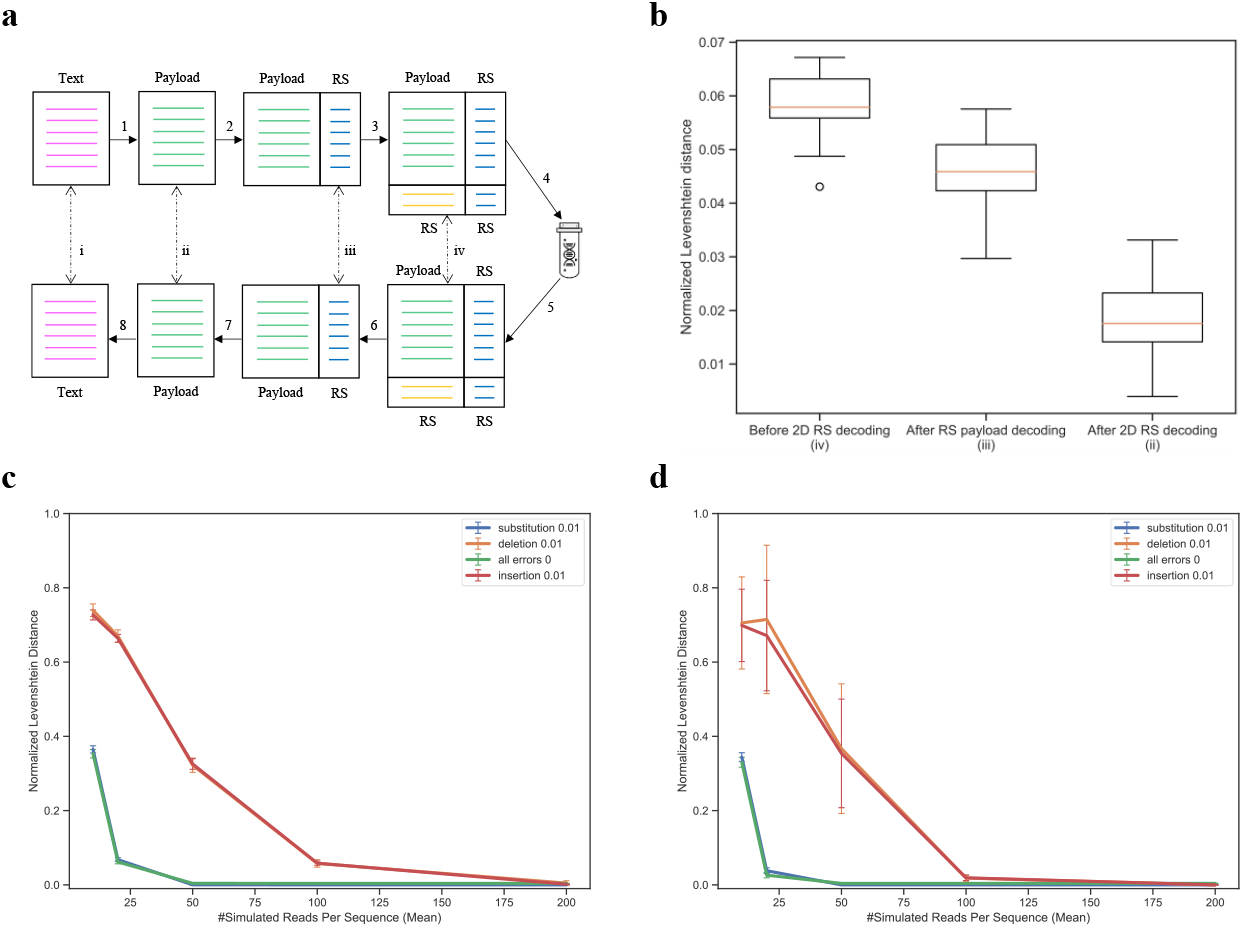
Simulation of an end-to-end combinatorial shortmer encoding. **a**, A schematic view of the simulation workflow. A text message is translated into a combinatorial message (1), and encoded using RS error correction on the barcode and payload (2). Each block is encoded using outer RS error correction (3). DNA synthesis and sequencing are simulated under various error schemes, and the combinatorial letters are reconstructed (4-5). RS decoding is performed on each block (6) and each sequence (7) before translation back to text (8). The Roman numerals (i-iv) represent the different error calculations. **b**, Error rates in different stages of the decoding process. Boxplot of the normalized Levenshtein distance (See Section 5.2.4) for the different stages in a simulation (30 runs) of sampling 100 reads, with an insertion error rate of 0.01. The X-axis represents the stages of error correction (before 2D RS decoding (iv), after RS payload decoding (iii), and after 2D RS decoding (ii)). **c, and d**, Sampling rate effect on overall performance. Normalized Levenshtein distance as a function of sampling rate before RS decoding (c) and after 2d RS decoding (ii). Different lines represent different error types (substitution, deletion, and insertion) introduced at a rate of 0.01.

**Fig. 4:**
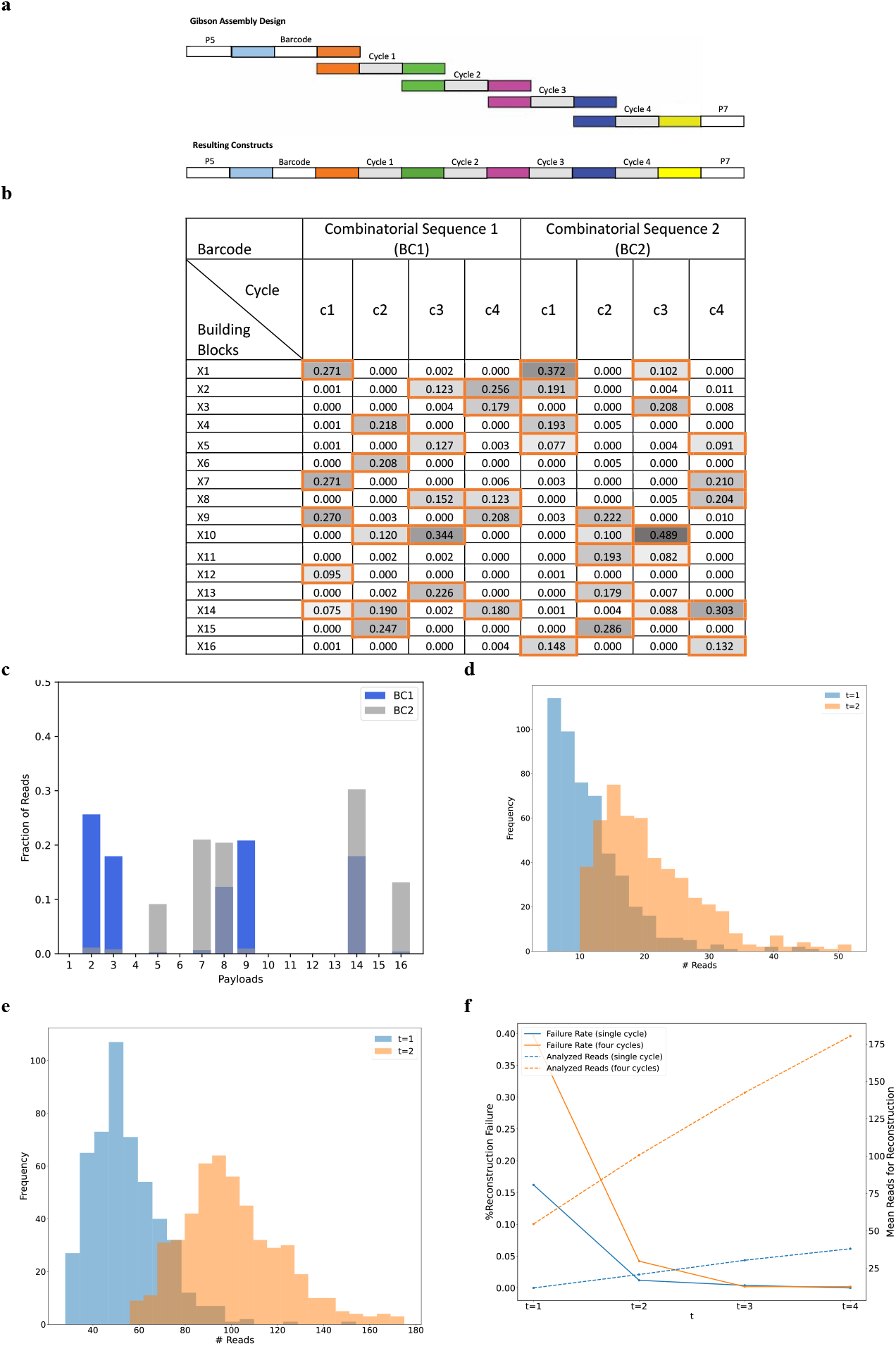
Experiment analysis. **a**, A schematic view of the Gibson assemby. Each combinatorial sequence consists of a barcode segment and four payload segments (denoted as cycle 1-4). **b**, Reconstruction results of the two combinatorial sequences. The color indicates read frequency and the member k-mers are marked with orange boxes. **c**, The distribution of reads over the 16 k-mers in an example combinatorial letter. Overlaid histograms represent the percentage of reads for each of the 16 k-mers for the same position in our two combinatorial sequences. This in fact, is an enlarged view of the two c4 columns of panel b. **d**, Required number of reads for reconstructing a single combinatorial letter. A histogram of the number of reads required to observe at least ***t*** = **1, 2** reads from ***K*** = **5** inferred k-mers. The results are based on resampling the reads 500 times, the data represents cycle 4. **e**, Required number of reads for reconstructing a four letter combinatorial sequence. Similar to d. **f**, Reconstruction failure rate as a function of the required multiplicity ***t***. Errornous reconstruction rate shown for different values of required copies to observe each inferred k-mer (***t*** = **1, 2, 3, 4**). The mean required number of reads for reconstruction is displayed using a secondary Y-axis in the dashed lines.

The results of the simulation runs are summarized in Fig. 3b-d. Each run included 30 repeats with random input texts of 10KB encoded using 98 combinatorial sequences, each composed of 134 combinatorial letters and 16nt barcode, as described above. Each run simulated the synthesis of 1000 molecules on average per combinatorial sequence and sampling of a subset of these molecules to be sequenced. The subset size was drawn randomly from *N*(*μ, σ* = 100), where *μ* is a parameter. Errors in predetermined rates were introduced during the simulation of both DNA synthesis and sequencing, as expected in actual usage [21] (See Section 5.2.3 for details on the simulation runs). Reconstruction rates and Levenshtein distances are calculated throughout the simulation process, as described in Fig. 3a.

Notably, the sampling rate is the dominant factor where even with zero synthesis and sequencing errors, low sampling rates yield such poor results (Fig. 3c) that the RS error correction is not able to overcome (Fig. 3d). The effect of substitution errors on the overall performance is smaller and they are also easier to detect and correct. This is because substitution errors occur at the nucleotide level rather than at the trimer level. The minimal Hamming distance *d* = 2 of the trimer set *Ω* allows for the correction of single-base substitutions. The use of 2D RS error correction significantly improved reconstruction rates, as can be observed in Fig. 3b.

### 3.5 Experimental proof of concept

To assess and establish the potential of large combinatorial alphabets, we also performed a small-scale experimental proof of concept. Gibson assembly was used to construct two combinatorial sequences, each containing a barcode and four payload cycles over a binomial alphabet with *N* = 16 and *K* = 5. The assembly was performed using DNA fragments composed of a 20-mer information sequence and an overlap of 20 bp between adjacent fragments, as shown in Fig. 4a. The assembled DNA was then stored and sequenced for analysis using Illumina Miseq (See Table 2 and Section 5.3 for details about the sequencing procedures). The sequencing output was then analyzed using the procedure described in Section 5.3.2. Both combinatorial sequences were successfully reconstructed from the sequencing reads, as presented in Fig. 4b and Supplementary Fig. 6, Fig. 7, and Fig. 8. The experiment also demonstrated the robustness of the binomial DNA encoding for synthesis and sequencing errors, as described in Fig. 4c. We observed a minor leakage between the two synthesized sequences, which was overcome by the reconstruction pipeline (See Fig. 4c and Supplementary Fig. 6, Fig. 7, and Fig. 8). Note that there is an overlap between the member k-mers of the two sequences.

**Table 2:**
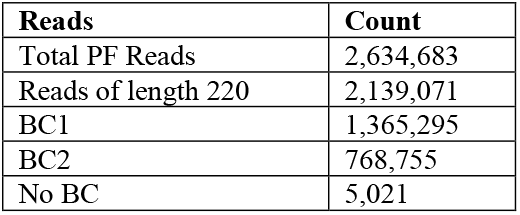
Summary of sequencing reads analyzed in the study. The table shows the total number of reads obtained, the number filtered by length (220 bases) for analysis, and the counts of reads associated with BC1, BC2, and those that did not have any recognizable barcode (No BC).

| Reads | Count |
| --- | --- |
| Total PF Reads | 2,634,683 |
| Reads of length 220 | 2,139,071 |
| BC1 | 1,365,295 |
| BC2 | 768,755 |
| No BC | 5,021 |

To test the effect of random sampling on the reconstruction of combinatorial sequences, we performed a subsampling experiment with *N* = 500 repeats, presented in Fig. 4d-f. We subsampled varying numbers of reads from the overall read pool and ran the reconstruction pipeline. Note that, as explained, the reconstruction of a single binomial position requires finding *K* = 5 inferred k-mers. That is, observing five unique k-mers at least *t* times. We tested the reconstruction performance using *t* = 1,2,3,4 and recorded the effect on the successful reconstruction rate and required number of reads.

For *t* = 1, reconstruction required analyzing 12.26 reads on average. These included 0.45 reads that contained an erroneous sequence that could not be mapped to a valid k-mer, and thus ignored. Note that the design of the set *Ω* of valid k-mers allows us to ignore only the reads for which the Hamming distance for a valid k-mer exceeded a predefined threshold (*d* = 3). If we ignored all the reads containing a sequence with non-zero Hamming distance to all k-mers, we would have skipped 2.26 extra reads, on average.

As expected, requiring *t* = 2 copies of each inferred k-mer resulted in an increase in the overall number of analyzed reads. Reconstruction of a single combinatorial letter required analyzing an average of 21.6 reads with 0.83 skipped and 3.99 non-zero Hamming distance reads. The complete distribution of the number of reads required for reconstruction of a single position using *t* = 1,2 is presented as a histogram in Fig. 4d.

To reconstruct a complete combinatorial sequence of 4 positions, we required the condition to hold for all positions. For *t* = 1, this entailed the analysis of 55.60 reads on average, out of which 1.04 reads were identified as erroneous and thus ignored, and with 7.36 non-zero Hamming distance reads. For *t* = 2, an average of 102.66 reads were analyzed with 1.97 skipped and 13.24 non-zero Hamming distance reads. The complete distribution of the number of reads required for reconstructing a complete combinatorial sequence using *t* = 1,2 is presented as a histogram in Fig. 4e.

Note that these results correspond to the analysis presented in Section 3.3, for the reconstruction of a single binomial position and a complete binomial sequence. Calculating the bound presented in Table 4, with *K* = 5 and *l* = 4, yields a requirement of approximately 140 reads to obtain 1 − *δ* = 0.99 probability of reconstruction. Clearly, this is well above the observed number of 55.60 reads. Note, as explained, the calculated bound is a loose bound. The reconstruction procedure ends with a set of inferred k-mers that represent the inferred combinatorial letter. This set is not guaranteed to be correct, especially when using *t* = 1, which means that noisy reads may result in an incorrect k-mer included in the inferred letter. Fig. 4f depicts the rate of incorrect reconstructions as a function of the number of required copies for each inferred k-mer (*t* = 1,2,3,4). Note that with *t* ≥ 3 results in 100% successful reconstruction. This, however, comes with a price, where more reads must be analyzed.

## 4 Discussion

In this study we introduced combinatorial shortmer encoding for DNA-based data storage, which extends the approach of composite DNA by addressing its key challenges. Combinatorial shortmer encoding allows for increased logical density, while ensuring low error rates and high reconstruction rates. We explored two encoding schemes, binary and binomial, and evaluated some of their theoretical and practical characteristics. The inherent consistency of the binomial encoding scheme, where every letter in the sequence consists of exactly K distinct member k-mers, ensures uniformity in the encoded DNA sequences. This approach not only simplifies the reading process, but also allows for a more streamlined decoding. For instance, technologies like nanopore sequencing enable continuous sequencing until all k-mers at a given position are confidently observed. On the other hand, the complexities introduced by binary encoding, which can yield a variable number of k-mers at any position, represent a potential challenge.

Similar to other DNA-based data storage systems, errors introduced to the sequences during the chemical and molecular stages affect the system’s performance. Our suggested approach is designed to inherently overcome base substitution errors, which are the most common errors expected in every DNA-based data storage system that includes DNA sequencing. This is achieved by the selection of a set of k-mers which is resilient to single-base substitutions, reducing the chances of letter mix-ups. Insertion and deletion errors, which usually originate in the synthesis process, are more challenging to overcome. We introduced a 2D RS error correction scheme on the shortmer level, allowing for a successful message reconstruction even with error levels exceeding those expected in reality.

Our study highlights the significant effect of sampling rates on the overall performance of the system. The accuracy and completeness of sequence reconstruction are closely tied to the rate at which DNA sequences are sampled. Optimal sampling rates ensure that the diverse regions of the encoded DNA are sufficiently represented, facilitating accurate reconstruction. An insufficient sampling rate can lead to data gaps, which further complicate the reconstruction process and may lead to errors or incomplete data retrieval. Our subsampling experiments underpin this observation, underscoring the need for calibration of sampling rates to ensure the desired fidelity in DNA-based data storage and retrieval.

While our proof-of-concept experiment showed success on a small scale, there are complexities to be addressed in considering large-scale applications. These include synthesis efficiency, error correction, and decoding efficiency. Nonetheless, the resilience of our binomial DNA encoding for both synthesis and sequencing errors highlights its practical potential and scalability.

Several future directions emerge from our study. First, it is essential to advance our error correction methods for better handling insertion and deletion errors. One approach for achieving this, is to adjust sampling rates: optimizing the sampling rate, especially in large-scale experiments, can lead to data retrieval at high accuracy. While our study highlighted the role of sampling rates in achieving desired outcomes, delve deeper into the underlying theory is necessary. By understanding the theoretical bounds of sampling rates, more concrete recommendations can be provided for real-world applications. Future research can further expand on this, both by conducting a series of experiments with varied sampling rates and by aiming to define theoretical bounds for these rates. This dual approach—combining practical experiments with rigorous theoretical analysis—could yield more precise guidelines for DNA-based data storage endeavors. Another future research direction, can be the development of error correction codes designed specifically to overcome the error types that characterize combinatorial encoding. Furthermore, transitioning from small-scale proof-of-concept experiments to larger-scale implementations is an important next step. Evaluating the scalability of our method across various scales and complexities will be enlightening, especially when considering synthesis efficiency and error rates. Finally, the consideration of advanced sequencing technologies could redefine the potential and efficacy of our proposed method.

## 5 Methods

### 5.1 Reconstruction probability of a binomial encoding letter

Let the number of reads required for reconstruction be a random variable 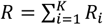 where *R*_1_ = 1 and 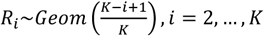 . Hence, the expected number of required reads, is:

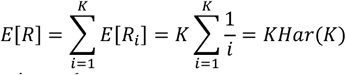

where *Har*(⋅) is the harmonic number.

Using the independence of *R*_*i*_, the variance of *R* can be bound by (See [22]):

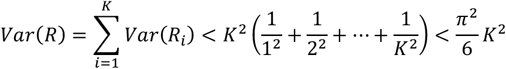

By Chebyshev’s inequality, we get an upper bound (a loose bound) on the probability of requiring more than *E*[*R*] + *cK* reads to observe at least one read of each member k-mer:

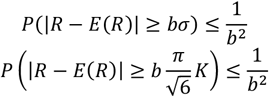

Let 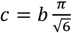, or 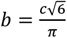 and we obtain:

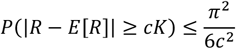

Or specifically

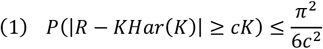

We now turn to address the reconstruction of an entire oligo of length *l*. Let *R*(*l*) be the random variable representing the number of reads required to have seen all the *K* member k-mers in every position. Setting any *δ* > 0, if we show that *P*(*R*(*l*) > *m*) ≥ 1 − *δ*, then we know that by accumulating *m* reads the probability of correct full reconstruction is more than 1 − *δ*. From equation (1), and assuming independence of the positions (in terms of observing all *K* member k-mers), we get equation (2):

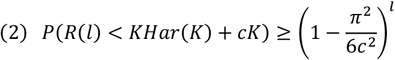

From which we can extract *c*, so that:

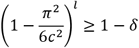

Which yields:

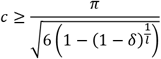

This process allows us to evaluate the sequencing depth complexity. For example, consider *l* = 100 and *δ* = 0.01. We want to find *c*, so that using *KHar*(*K*) + *cK* reads will reconstruct the entire sequence with 0.99 probability. We therefore set:

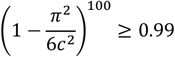

And get:

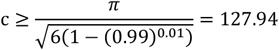

And therefore, using 128 reads guarantees reconstruction with 0.99 probability.

### 5.2 An end-to-end combinatorial storage system

In section 3.4 we propose an end-to-end combinatorial storage system, as follows.

#### 5.2.1 Combinatorial encoding and padding

A binary message is encoded using a large k-mer combinatorial alphabet (e.g., trimer-based alphabet of size |*∑*| = 4096 letters, with *N* = |*Ω*| = 16), resulting in *r* = 12 bits per combinatorial letter. The binary message is zero padded to ensure its length is divisible by *r* prior to the combinatorial encoding. The complete message is broken into sequences of set length *l* = 120, each sequence is then marked with a standard DNA barcode and translated using the table presented in the Encode legend (See Section 7.2).

The length of the complete combinatorial sequence must be divisible by the payload size *l* and by the block size *B*. As described in Fig. 5, this is ensured using another padding step, and the padding information is included in the final combinatorial sequence.

**Fig. 5:**
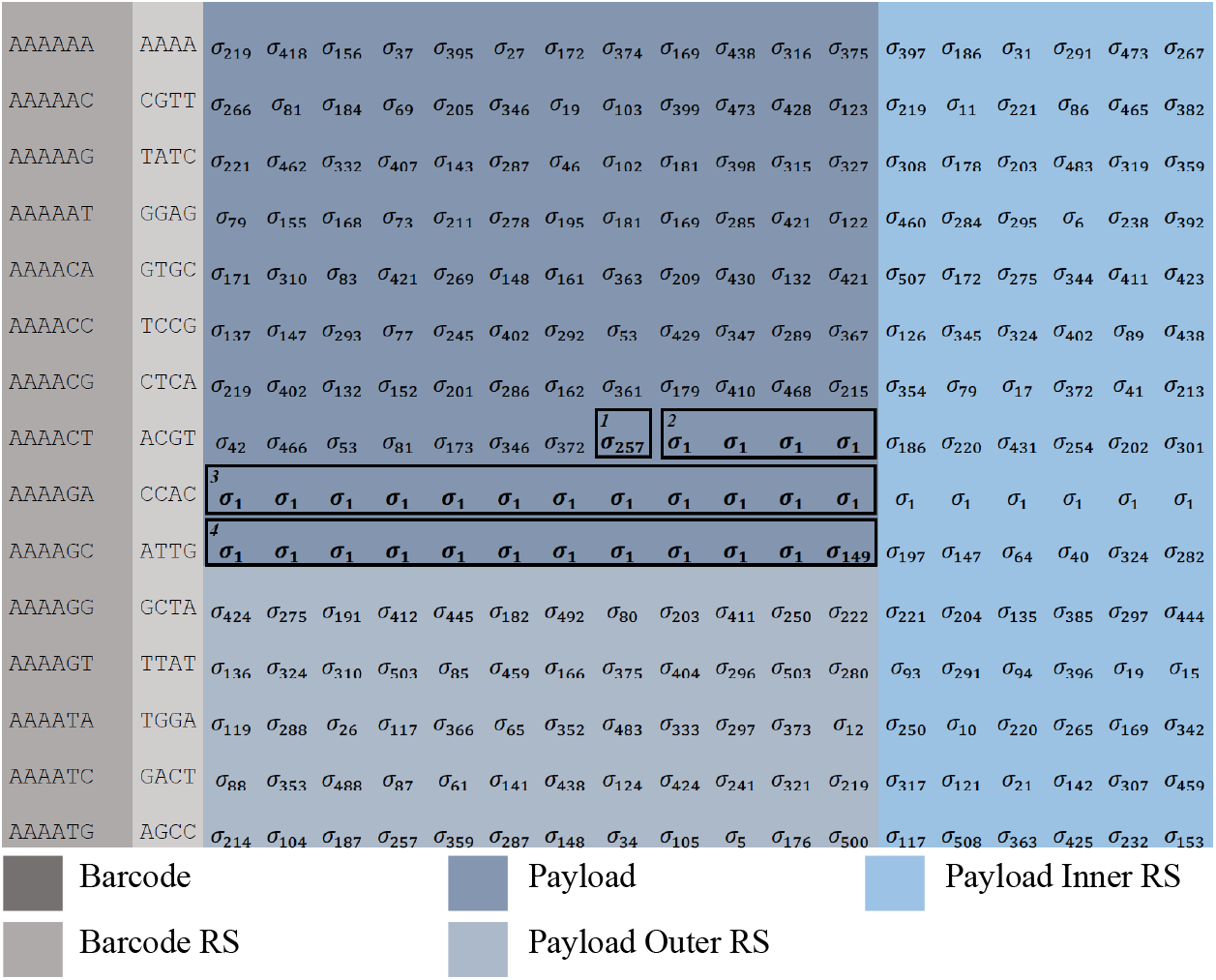
Example of message coding including padding and Reed-Solomon error correction. Encoding of a ∼0.1KB message to a 512 letter binomial alphabet (N= **16, *K*** = **3**). First, bit padding is added, included here in the letter 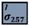 . Next, block padding is added, included here in 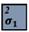 and 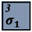 . Padding information is included in the last sequence of all blocks. The last sequence holds the number of padding binary bits. In this example, 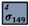 represents 148 bits of padding, composed of **4** + (**4** ∗ **9**) + (**12** ∗ **9**) ***bits***, 4 bits from 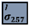, 4 letters from 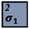 and 12 letters from 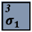 .

#### 5.2.2 Error correction codes

The two-dimensional (2D) error correction scheme includes using three Reed Solomon (RS)

[23] encodings: on each barcode, on the payload part of each sequence, and an outer error correction code on each block of sequences.

- Each barcode is encoded using a systematic RS(6,8) code over *GF*(2^4^), transforming the unique 12nt barcode into a 16nt sequence.
- Each 120 combinatorial letter payload sequence is encoded using a RS(120,134) code over *GF*(2^12^), resulting in a sequence of length 134 combinatorial letters.
- To protect against sequence dropouts, outer error correction code is used on the columns of the matrix (See Fig. 5). Each block of *B* = 42 sequences, is encoded using a RS(42,48) RS code *GF*(2^12^). This is applied in each column separately.

For simplicity, Fig. 5 demonstrates the encoding of ∼0.1 KB using shorter messages with simpler error correction codes. The following parameters are used:

- A barcode length of 6nt encoded using RS(3,5) code over *GF*(2^4^) to get 10nt.
- A payload length of *l* = 12 encoded using RS(12,18) over *GF*(2^9^) for the alphabet. 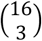 binomial
- A 10-sequence block encoded, column wise, using a (10,15) RS code over *GF*(2^9^). The 824 bits are first padded to be 828 = 92 ∗ 9. The 92 combinatorial letter message is split into 7 sequences of 12 letters and an additional sequence of 8 letters. Finally, a complete block of 12 sequences (total of 10 ∗ 12 = 120 letters) is created by padding with one additional sequence of 12 letters and including the padding information as the last sequence.

#### 5.2.3 Synthesis and sequencing simulation with errors

- **Simulating the synthesis process**. DNA molecules pertaining to the designed sequences are synthesized using combinatorial k-mer DNA synthesis (See Fig. 1b). For each combinatorial sequence, we first determine the number of synthesized copies by sampling from *X*∼*N*(*μ* = 1000, *σ*^2^ = 100). Let *x* be the number of copies for a specific sequence. Next, for every position in the sequence we uniformly sample *x* independent k-mers from the set of member k-mers of the combinatorial letter in the specific position. We concatenate the sampled k-mers to the already existing *x* synthesized molecules.
- **Error simulation**. Synthesis and sequencing error are simulated as follows. Error probabilities for deletion, insertion, and substitution are given as parameters denoted as *P*_*d*_, *P*_*I*_, and *P*_*s*_ respectively. Deletion and Insertion errors are assumed to occur during k-mer synthesis and thus implemented on the k-mer level (i.e., an entire k-mer is deleted or inserted in a specific position during the synthesis simulation). Substitution errors are assumed to be sequencing errors, and hence implemented on a single base level (i.e., a single letter is substituted, disregarding the position within the k-mer).
- **Mixing**. Post synthesis, molecules undergo mixing to mirror genuine molecular combinations. This is achieved through a randomized data line shuffle using a SQLite database, enabling shuffle processes even for sizable input files [24].
- **Reading and sampling**. From the simulated synthesized molecule set, a subsample of predefined size *S* ∗ *number of synthesized seqeunces* is drawn, simulating the sampling effect of the sequencing process.

#### 5.2.4 Reconstruction

- **Barcode decoding**. The barcode sequence of each read is decoded using the RS(6,8) code.
- **Grouping by barcode**. The reads are then grouped by their barcode sequence to allow the reconstruction of the combinatorial sequences.
- **Filtering of read groups**. Barcodes (set of reads) with less than 10% of the sampling rate *S* reads are discarded.
- **Combinatorial reconstruction**. For each set of reads, every position is analyzed separately. The *K* most common k-mers are identified and used to determine the combinatorial letter *σ* in this position. Let Δ be the difference between the length of the analyzed reads and the length of the designed sequence. Δ = *l* − *len*(*read*). Reads with |Δ| > *k* − 1 are discarded from the analysis. Invalid k-mers (not in *Ω*) are replaced by a dummy k-mer *X*_*dummy*_.
- **Missing barcodes**. Missing barcodes are replaced with dummy sequences to enable correct outer RS decoding.
- **Normalized Levenshtein distance**. Levenshtein distance between the observed sequence O and the expected sequence *E* is calculated [25] [26]. Normalized Levenshtein distance is calculated by dividing the distance by the length of the expected sequence:

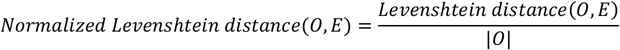

### 5.3 Proof of concept experiment

The proof-of-concept experiment was performed by imitating combinatorial synthesis using Gibson assembly of larger DNA fragments. Each DNA fragment was composed of a 20-mer information sequence and an overlap of 20 bp between adjacent fragments, as depicted in Fig. 4a. Two combinatorial sequences were designed, each composed of a barcode fragment, 4 payload fragments, and Illumina P5 and P7 anchors at the ends. The information fragments included in each combinatorial position were chosen from a set of 16 sequences with sufficient pair-wise distance. The full list of DNA sequences and the design of combinatorial sequences is listed in Supplementary Section 7.5.

No animal, or human participants were involved in the study.

#### 5.3.1 DNA assembly and sequencing

Payload, barcode, and P7 anchor fragments with 20 bp overlaps for the purpose of Gibson assembly were produced by annealing complementary oligonucleotides manufactured by Integrated DNA Technologies (IDT). Oligos were dissolved in Duplex Buffer (100 mM Potassium Acetate; 30 mM HEPES, pH 7.5; available from IDT) to the final concentration of 100 micromolar. For annealing, 25 microliters of each oligo in a pair were combined to the final concentration of 50 micromolar. The oligo mixes were incubated for 2 min at 94^0^ C and gradually cooled down to room temperature. The annealed payload oligos that belonged to the same cycle (5 oligos total) were mixed to the final concentration of 1 micromolar per oligo – a total of 5 micromolar, by adding 2 microliters of each annealed oligo into the 90 microliters of nuclease-free water – a final volume of 100 microliters. Annealed barcode and P7 anchor oligos were also diluted to the final concentration of 5 micromolar in nuclease-free water, after thorough mixing by vortexing. The diluted oligos were stored at -20^0^C.

Immediately prior to the Gibson assembly, payload oligo mixes, barcode, and P7 anchor oligos were further diluted 100-fold to the final working dilution of 0.05 pmol/microliter in nuclease-free water. Gibson reaction was assembled by adding 1 microliter (0.05 pmol) of barcode, 4 x cycle mixes, and P7 anchor to the 4 microliters of nuclease-free water and supplemented with 10 microliters of NEBuilder HiFi DNA assembly master mix (New England Biolabs (NEB)) to the final volume of 20 microliters according to the manufacturer instructions. The reactions were incubated for 1 hr at 50^0^C and purified with AmpPure Beads (Thermo Scientific) at 0.8X ratio (16 microliters of beads per 20 microliters Gibson reaction) to remove free oligos / incomplete assembly products. After adding beads and thorough mixing, the reactions were incubated for 10 min at room temperature and then placed on a magnet for 5 min at room temperature. After removing the sup, the beads were washed twice with 100 microliters of 80% ethanol. The remaining washing solution was further removed by a 20 microliter tip and the beads dried for 3 min on the magnet with an open lid. After removing from the magnet, the beads were resuspended in 22 microliters of IDTE buffer (IDT), incubated for 5 min at room temperature, and then placed back on the magnet.

20 microliter of eluate were transferred into the separate 1.7 ml tube. 5 microliters of the eluted DNA were used as a template for PCR amplification combined with 23 microliters of nuclease-free water, 1 microliter of 20 micromolar indexing primer 5, 1 microliter of 20 micromolar indexing primer 7, and 10 microliters of rhAMPseq master mix v8.1 – a total of 40 microliters. After initial denaturation of 3 min at 95^0^C, the PCR reaction proceeded with 50 cycles of 15 sec at 95^0^C, 30 sec at 60^0^C, and 30 sec at 72^0^C, followed by final elongation of 1 min at 72^0^C and hold at 4^0^C. The PCR reactions were purified with Ampure beads at 0.8X ratio (32 microliter beads per 40 microliters of PCR reaction) as outlined above and eluted in 22 microliters IDTE buffer. The concentration and the average size of the eluted product were determined by Qubit High Sensitivity DNA kit and Agilent 2200 TapeStation system with D1000 high-sensitivity screen tape respectively. The eluted product was diluted to 4 nanomolar concentration and used as an input for denatured sequencing library preparation, per manufacturer instructions. The sequencing was performed on Illumina Miseq apparatus (V2 chemistry, 2 x 150 bp reads) using 6 picomolar denatured library supplemented with 40% PhiX sequencing control.

#### 5.3.2 Decoding and analysis

This section outlines the key steps involved in our sequencing analysis pipeline, aimed at effectively processing and interpreting sequenced reads. The analysis pipeline gets the sequencing output file containing raw reads in “.fastq” format and a design file containing the combinatorial sequences.

Analysis Steps:

1. **Length Filtering**. We saved reads that were 220 bp in length, retaining only those corresponding to our designed read length.
2. **Read Retrieval**. We carefully checked each read for the presence of BCs, universals, and payloads. To keep our data accurate, we discarded reads where the BCs, universals, or payloads had a Hamming distance of more than 3 errors.
3. **Identifying inferred k-mers**. For every BC and each cycle, we counted the K most common k-mers. We then compared these with the design file to countify those matching (Fig. 4b).

### 5.4 Information capacities for selected encodings

Table 1 illustrates the logical densities derived from encoding a 1 GB binary message using oligonucleotides with a 12nt barcode and an additional 4nt for standard DNA Reed-Solomon (RS) error correction, and a 120 letters payload with 14 extra RS for the payload in combinatorial encoding schemes with parameters N and K.

The densities were calculated as follows:

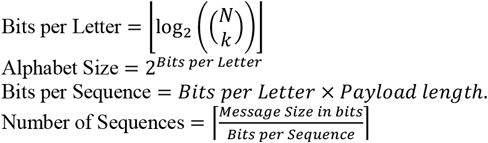

Number of sequences padded: Total number of sequences after padding for the block size.

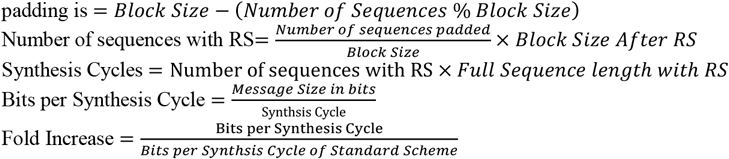

## Supporting information

Combinatorial shortmer synthesis (video)

Bionomial shortmer alphabet example (Table)

Proof-of-concept experimental design (Table)

## 7 Supplementary

All files are available [here](https://drive.google.com/drive/folders/15k8-5Cgq_BPjazBVv4-n6vBzyIDd7kIb?usp=drive_link)

### 7.1 Combinatorial shortmer synthesis (video)

Supplementary video, uploaded separately in:

Supplementary_animation_3-mers_16choose2.wmv.

### 7.2 Bionomial shortmer alphabet example (Table)

Supplementary file, uploaded separately in:

Supplementary_table_alphabet.xlsx

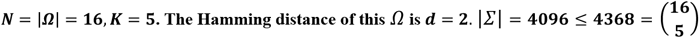

### 7.3 Example of k-mer sets

Table 3 is an example of k-mer sets. To the left, are two sets of trimers that have a minimal Hamming distance of 2, with |*Ω*_1_| = 16 *and* |*Ω*_2_| = 12. To the right, is a set of 54 6-mers that have a minimal Hamming distance of 4.

**Table 3:**
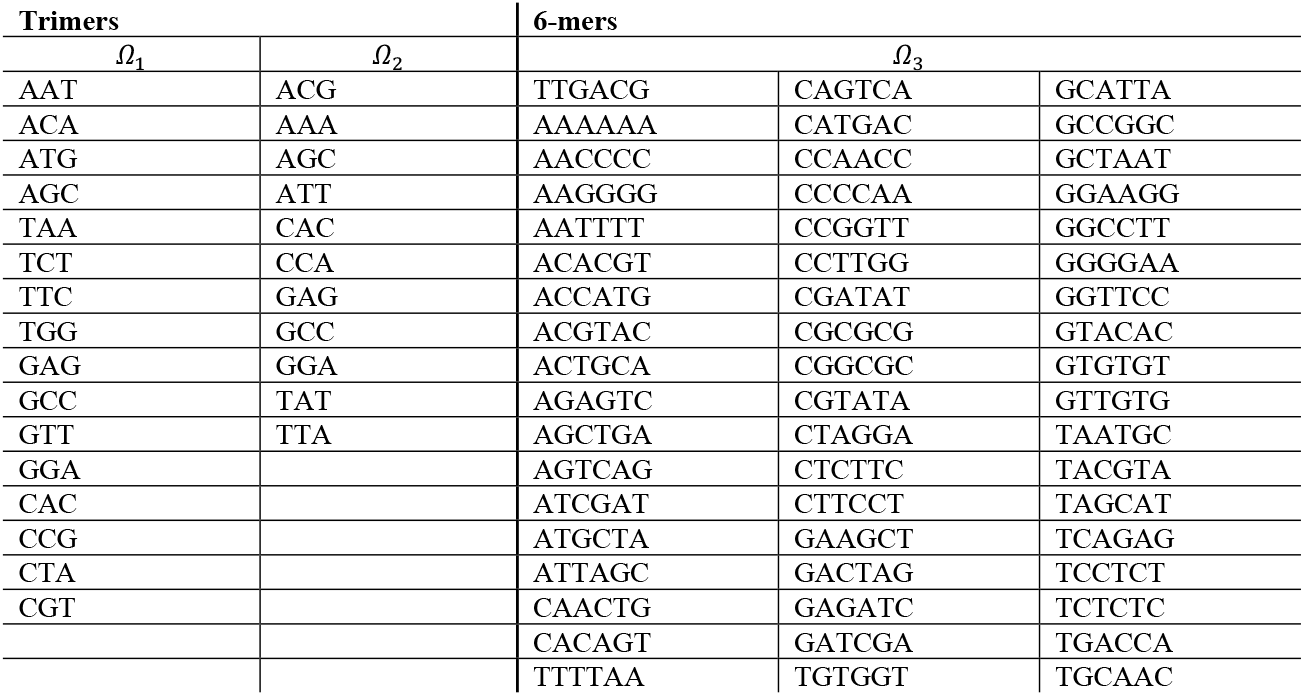
Example of k-mer sets, *Ω*. For ***Ω***_**1**_ and ***Ω***_**2**_, the minimum Hamming distance is 2. For ***Ω***_**3**_, the minimum hamming distance is 4.

### 7.4 Reconstruction of a binomial seuqence

**Table 4:**
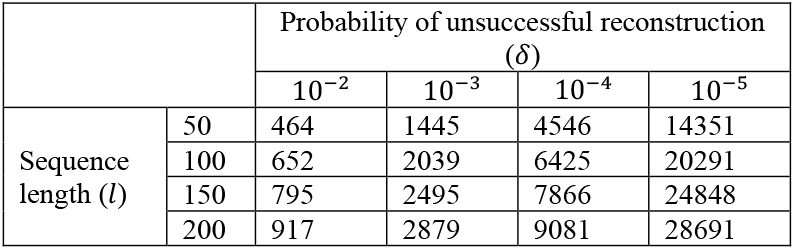
Sufficient number of reads to reconstruct a binomial sequence. Entries in the table represent ***KHar***(***K***) + ***cK*** where ***c*** is derived based on the desired ***δ***, as explained in Section 5.1.

### 7.5 Proof-of-concept experimental design (Table)

Supplementary file, uploaded separately in:

Supplementary_table_oligo_sequences.xlsx

### 7.6 Proof of concept smaller-scale experiment

For each of the two barcodes, we were able to identify and recover the barcode and their payloads. At each position/cycle in the sequence, the five-member k-mers were recovered, which are the five inferred k-mers (See Fig. 6 and Fig. 7).

**Fig. 6:**
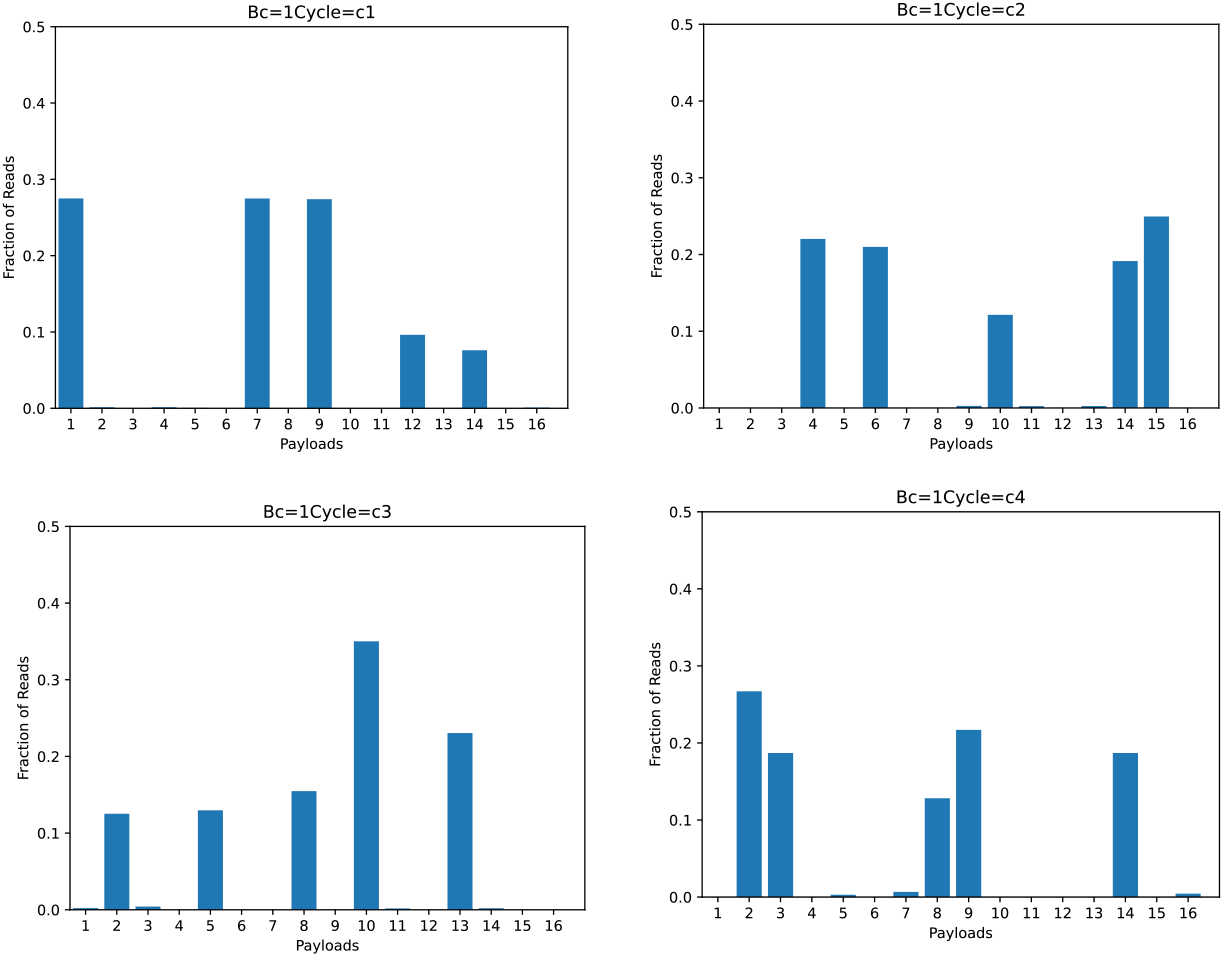
Smaller-scale experiment, oligo 1. Oligo has four cycles, each with five inferred k-mers. Results showed the k-mers expected, according to our original design.

**Fig. 7:**
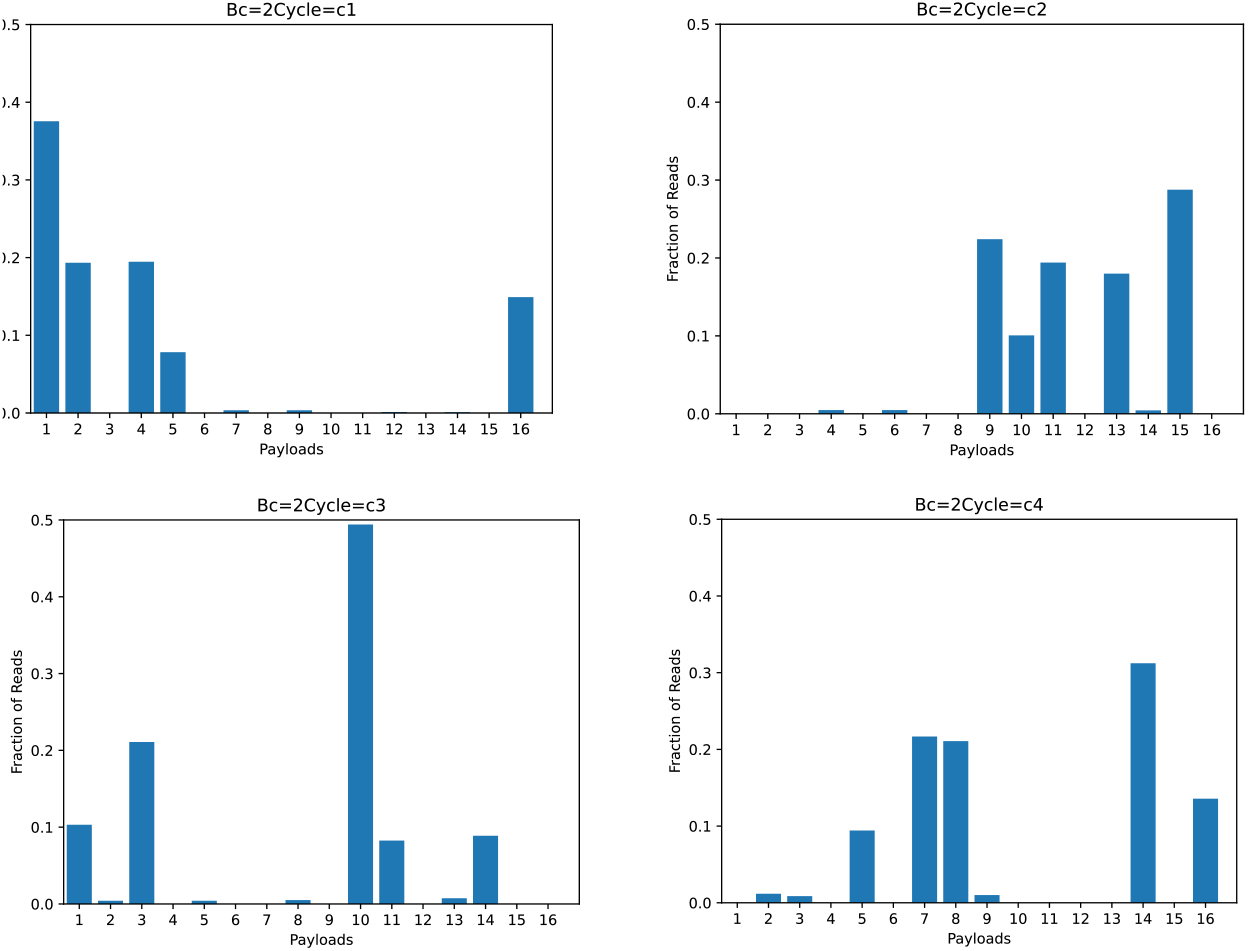
Smaller-scale experiment, oligo 2. Oligo has four cycles, each with five k-mers. Results showed the k-mers expected, according to our original design.

**Fig. 8:**
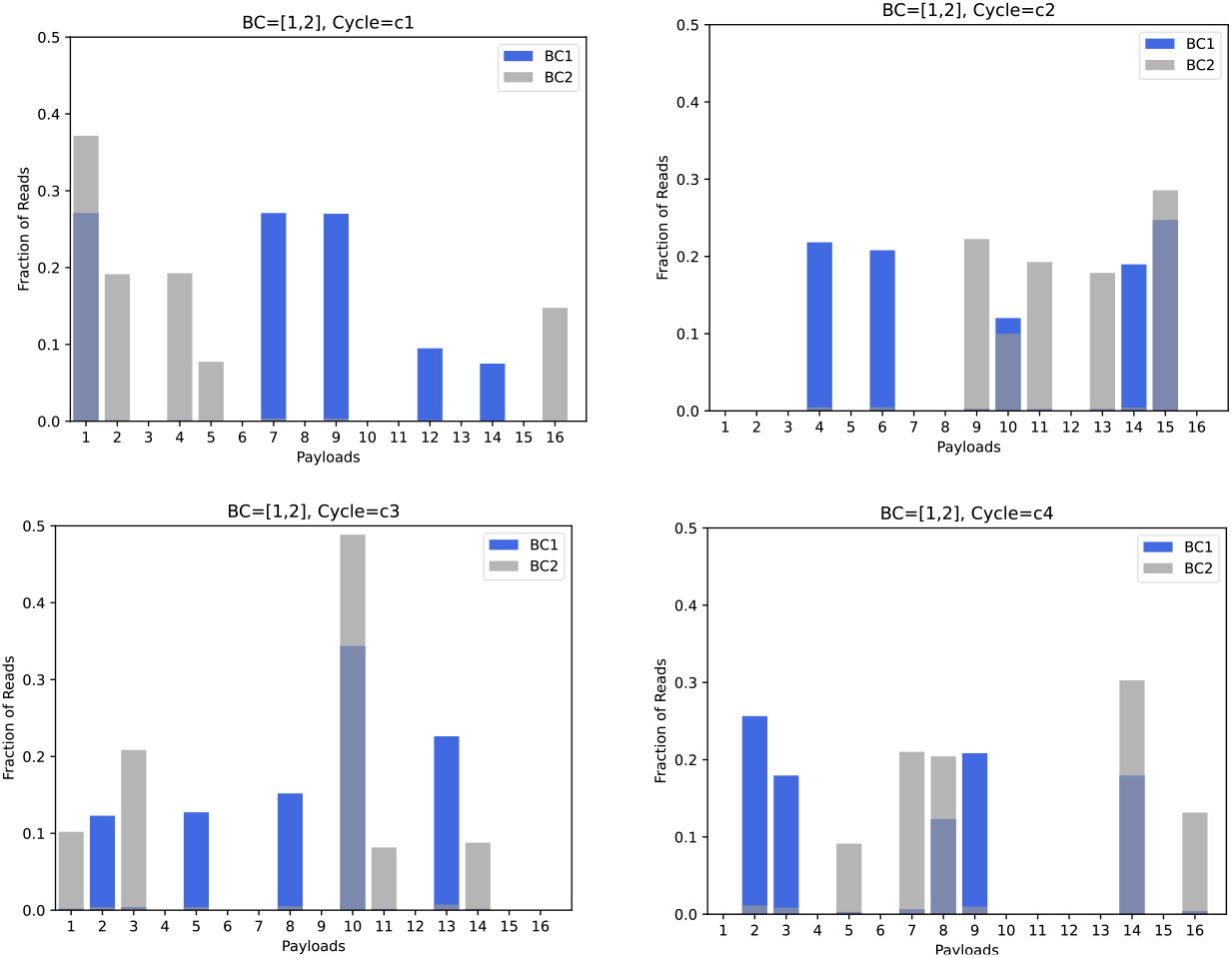
Superimposing oligo 2 (grey) over oligo 1 (blue). Leakage resulted in more than five k-mers expected, according to our design, where k-mers continued to assemble in each cycle due to the active enzyme, yet not necessarily on their designated oligo. The same applies to the second sequence that was assembled. Note that there is an overlap between the member k-mers of the two sequences. See for example, Payloads #10 and #15 in cycle C2.

## Notes

### Competing Interest Statement

Patent Ownership and Research Subject: Leon Anavy and Zohar Yakhini are listed as inventors on a patent that is directly related to the research presented in this submission. The details of the patent are as follows:
Title: "Molecular data storage systems and methods"
Patent Number: US20210141568A1
Year: 2021
Relationship to Submitted Work: The research presented in this submission is directly based on, or is an extension of, the concepts and technologies described in the patent.
The authors confirm that there are no other competing interests, financial or otherwise, that could be perceived to influence, or that give the appearance of potentially influencing, the work submitted.

### Summary of Updates

- Enhanced Clarity and Detail: Improved explanations and comprehensive details, particularly in methodology and results. - Updated Figures: Revised figures for clearer concept and result illustration. - Refined Analysis and Simulations: Updated analysis and simulation data for a more in-depth exploration of DNA storage methods. - Real DNA Experiments: Incorporation of real DNA experiments to demonstrate the proof of concept for the proposed storage method. - Cover Depth Analysis: New emphasis on the importance of cover depth analysis in combinatorial DNA-based storage systems. - Revised Author Affiliations: Updated author affiliations. - Updated Supplemental Material: Modified supplemental files to align with the main text revisions.

