## Supplementary Material for "Efficient DNA-based data storage using shortmer combinatorial encoding"

### 7 Supplementary

All files are available here

#### 7.1 Combinatorial shortmer synthesis (video)

Supplementary video, uploaded separately in:
Supplementary_animation_3-mers_16choose2.wmv.

#### 7.2 Bionomial shortmer alphabet example (Table)

Supplementary file, uploaded separately in:
Supplementary_table_alphabet.xlsx

**$N = |\Omega| = 16, K = 5$. The Hamming distance of this $\Omega$ is $d = 2$. $|\Sigma| = 4096 \leq 4368 = \binom{16}{5}$**

#### 7.3 Example of k-mer sets

Table 3 is an example of k-mer sets. To the left, are two sets of trimers that have a minimal Hamming distance of 2, with $|\Omega_1| = 16$ *and* $|\Omega_2| = 12$. To the right, is a set of 54 6-mers that have a minimal Hamming distance of 4.

| **Trimers** | | **6-mers** | | |
|---|---|---|---|---|
| $\Omega_1$ | $\Omega_2$ | $\Omega_3$ | | |
| AAT | ACG | TTGACG | CAGTCA | GCATTA |
| ACA | AAA | AAAAAA | CATGAC | GCCGGC |
| ATG | AGC | AACCCC | CCAACC | GCTAAT |
| AGC | ATT | AAGGGG | CCCCAA | GGAAGG |
| TAA | CAC | AATTTT | CCGGTT | GGCCTT |
| TCT | CCA | ACACGT | CCTTGG | GGGGAA |
| TTC | GAG | ACCATG | CGATAT | GGTTCC |
| TGG | GCC | ACGTAC | CGCGCG | GTACAC |
| GAG | GGA | ACTGCA | CGGCGC | GTGTGT |
| GCC | TAT | AGAGTC | CGTATA | GTTGTG |
| GTT | TTA | AGCTGA | CTAGGA | TAATGC |
| GGA | | AGTCAG | CTCTTC | TACGTA |
| CAC | | ATCGAT | CTTCCT | TAGCAT |
| CCG | | ATGCTA | GAAGCT | TCAGAG |
| CTA | | ATTAGC | GACTAG | TCCTCT |
| CGT | | CAACTG | GAGATC | TCTCTC |
| | | CACAGT | GATCGA | TGACCA |
| | | TTTTAA | TGTGGT | TGCAAC |

**Table 1: Example of k-mer sets, $\Omega$.** For $\Omega_1$ and $\Omega_2$, the minimum Hamming distance is 2. For $\Omega_3$, the minimum hamming distance is 4.

#### 7.4 Reconstruction of a binomial seuqence

| | | Probability of unsuccessful reconstruction ($\delta$) | | | |
|---|---|---|---|---|---|
| | | $10^{-2}$ | $10^{-3}$ | $10^{-4}$ | $10^{-5}$ |
| Sequence length ($l$) | 50 | 464 | 1445 | 4546 | 14351 |
| | 100 | 652 | 2039 | 6425 | 20291 |
| | 150 | 795 | 2495 | 7866 | 24848 |
| | 200 | 917 | 2879 | 9081 | 28691 |

**Table 2: Sufficient number of reads to reconstruct a binomial sequence.** Entries in the table represent $\boldsymbol{KHar(K) + cK}$ where $\boldsymbol{c}$ is derived based on the desired $\boldsymbol{\delta}$, as explained in Section **Error! Reference source not found.**.

#### 7.5 Proof-of-concept experimental design (Table)

Supplementary file, uploaded separately in: Supplementary_table_oligo_sequences.xlsx

#### 7.6 Proof of concept smaller-scale experiment

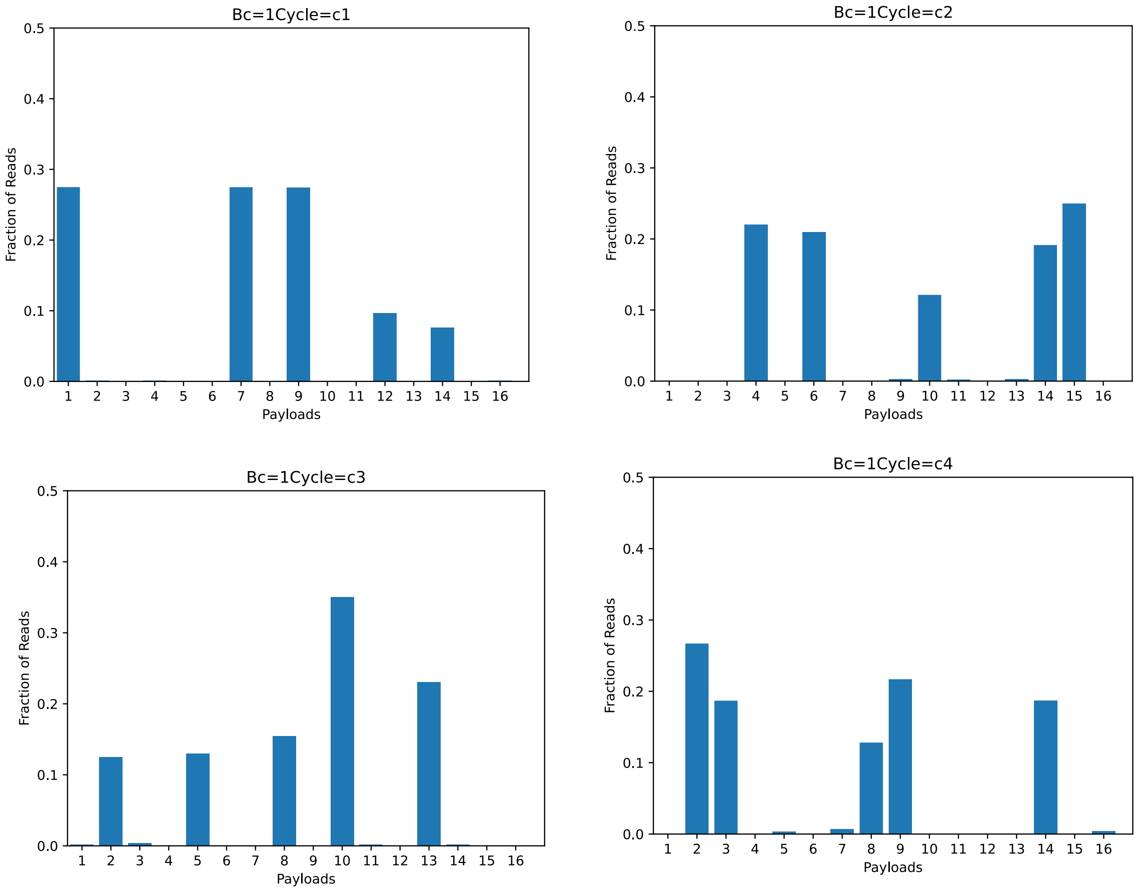


**Fig. 1: Smaller-scale experiment, oligo 1.** Oligo has four cycles, each with five inferred k-mers. Results showed the k-mers expected, according to our original design.

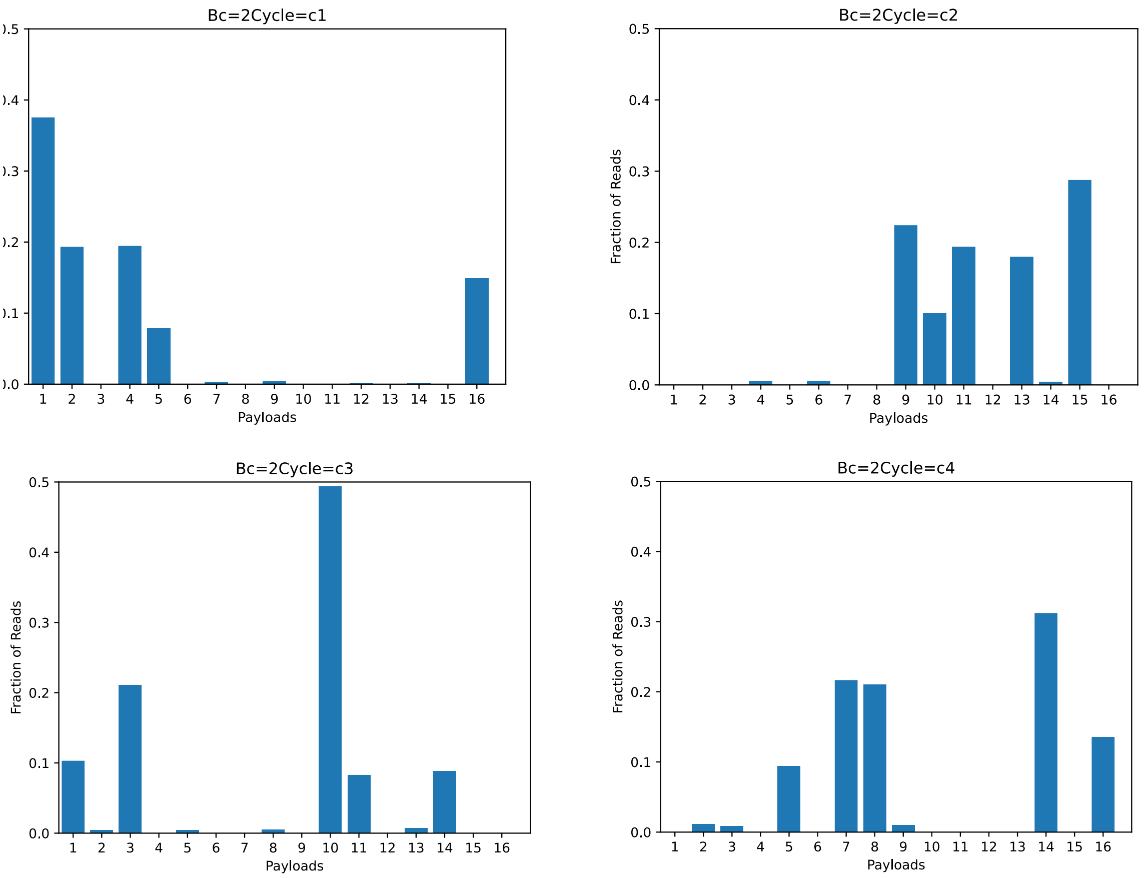


**Fig. 2: Smaller-scale experiment, oligo 2.** Oligo has four cycles, each with five k-mers. Results showed the k-mers expected, according to our original design.

For each of the two barcodes, we were able to identify and recover the barcode and their payloads. At each position/cycle in the sequence, the five-member k-mers were recovered, which are the five inferred k-mers (See Fig. 1 and Fig. 2).

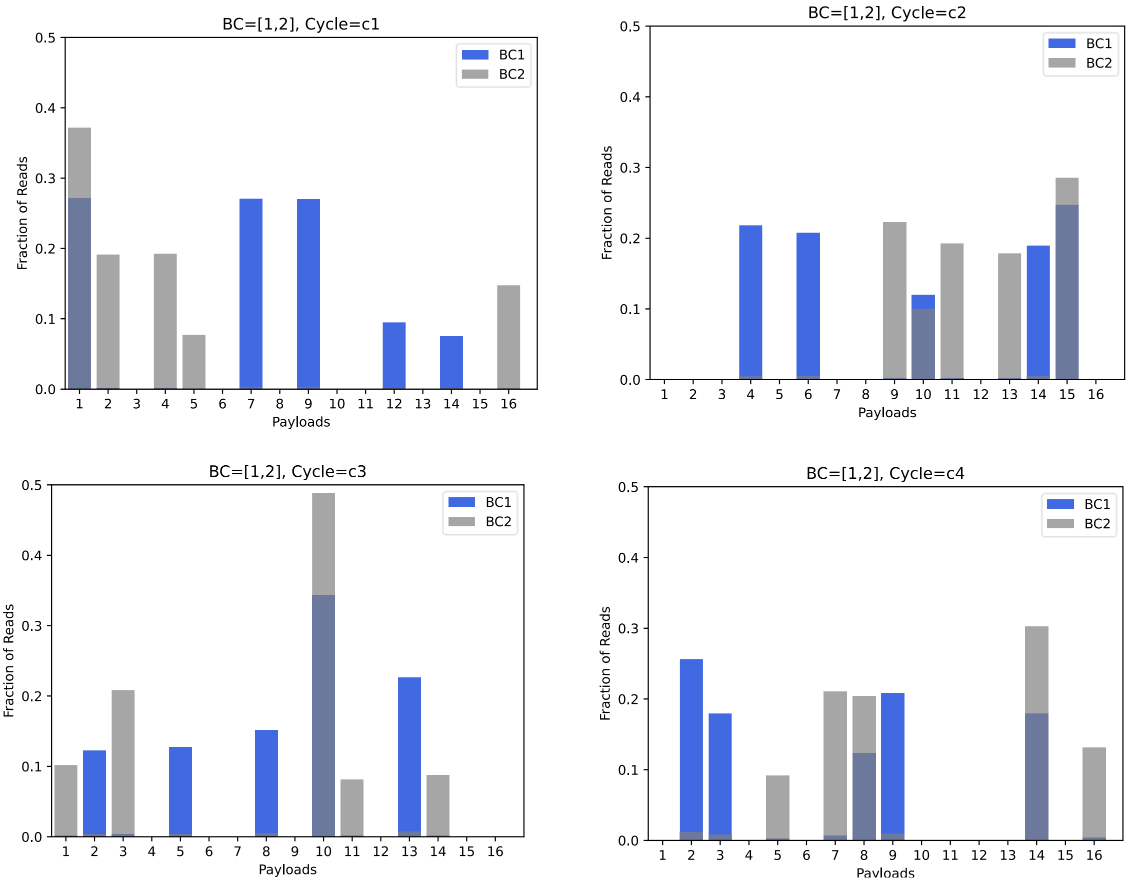


**Fig. 3: Superimposing oligo 2 (grey) over oligo 1 (blue).** Leakage resulted in more than five k-mers expected, according to our design, where k-mers continued to assemble in each cycle due to the active enzyme, yet not necessarily on their designated oligo. The same applies to the second sequence that was assembled. Note that there is an overlap between the member k-mers of the two sequences. See for example, Payloads #10 and #15 in cycle C2.
